# Molecular Dynamics Simulations of Phytopthora sojae Avirulence Factor 5, PsAvh5, Define Membrane Binding, Inositolphosphate-3’-phosphate Interactions and Protein Mechanics

**DOI:** 10.1101/2021.01.23.427770

**Authors:** Stephan L. Watkins

## Abstract

Phytopthora Avirulence proteins are a primary target for development of rational chemical and biological control of some of the most devastating plant pathogens. Despite the sequencing of entire genomes, and characterization of many of these proteins at the chemical level, many questions remain regarding actual chemical and biological interactions involved. In addition, disputed roles of ligands, such as Inositolphosphate-3’-phosphate and amino acids of important function remain unclear. To address some of these issues, we developed molecular models from structural elements and published data for Phytopthora sojae avirulence protein 5. Molecular dynamics simulations are used to study protein function, interactions involved primarily with lipids and membranes, and inositol derivatives. Our findings indicate that the protein is stable as a monomer, and in a dimeric form. Also, that these proteins interact with Inositolphosphate-3’phosphate as a necessary membrane element, in binding. We identified several amino acids of importance, additional to defining the mechanical features of the protein within the binding process to different membranes. A high affinity, comparable to other membrane surface binding molecules of −219.54 Kcal for the dimer, and −176.61 for the monomer were determined. With either form, we found the inositolphosphate-3’-phosphate to be essential in the membrane binding process. Our findings answer some of the debated questions while creating a point to further test avirulence proteins in general for functional aspects. Additionally, the structures and data can be utilized to provide a better starting point for rational design approaches to control this pathogen.

## Introduction

Potential models of avirulence protein (Avr) mechanics from Phytophthora pathogens is a primary focus for agricultural and forestry based industries, and many government agencies [1,2]. Working understanding of the functionality of these at the amino acid level provides the foundation for interventive measures. Phytophthora pathogens themselves are responsible for billions of dollars in food crop loss annually, and have now wiped out single species of trees in European forest at a 30-50% rate in some areas. In the US, P. ramorum has taken a toll on oak populations, especially in California [3]. In Europe the same pathogen reached as much as 50% mortality in Germany for unrelated tree species such as beech, maple and larch, and has infected maple and oak with high mortality across Switzerland [4,5,6]. Comparatively, the strains P. alni has now begun to infect alder trees, and P. infestans cyclically kills Solanaceae species, such as potato in Ireland [7]. Starting in 1840, P. infestans alone caused the well-noted Irish potato famine, reemerging every 30 to 40 years, the last wave being in 1980 [8,9]. Many infestations and forest or crop based damage are relatively new, due in part to globalized trade. Rapid spread threatens forest or food crops at an alarming rate, influencing national production from agriculture sectors and individual farms. Phytophthora infestans alone can destroy an entire agricultural region in as little as 10 days. These threats warrant immediate responses from government agencies or international sources to combat the effects to national and global security [10,11].

Methods for control of these pathogens vary widely. Traditionally copper compounds or ranges of amine based chemical agents have been used [12]. For this later, organisms seem to rapidly adapt resistance to the effects of the compounds, leaving only copper based compounds with higher environmental toxicity and allergy based reactions as a means of control [13,14]. In many countries, responses have been to utilize selective plant breeding based on resistant plant strains. This is problematic, as most agriculture based plants have lost resistance genes for specific Phytophthora strains in the course of domestication [15]. Native plants are often rare or nonexistent but necessary to re-introduce resistance. In many cases, as observed in Switzerland, Italy, Germany and France the only course of action has been complete destruction of all infected plants when they become parasitized. For forests, this has resulted in widespread loss of tree crops that require decades for regeneration, and has taken a toll on preserved forestlands.

Currently, understanding of protein structure and function has lagged behind larger whole genome projects, despite the availability of base genetic information [16]. Structural based information is necessary for rational and rapid design of new chemical, peptide or protein based methods in controlling Phytophthora [17]. Many basic questions remain, such as actual final amino acid sequences of proteins, secondary modifications, cellular location in either host or pathogen, and domain and amino acid interaction with specific ligands [18]. Many whole genomes have been sequenced allowing for whole genome cross species comparisons and extensive evolutionary studies [11,19]. Data intensive research has been conducted rapidly, such as yeast two hybrid screening of Phytophthora host plant interactions, host or pathogen RNA sequencing, and sequence data mining [16]. Conversely, basic experiments such as single protein affinities, protein posttranslational modifications, or understanding of Phytophthora cellular processing at the biochemical level have not been conducted. Only recently has whole genome comparisons raised the question of large differences in splicing, secretion or compartmental peptide sequence tags within these pathogenic organisms [20].

Ecological studies combined with genetics, have recently shifted the focus to proteins in describing the cyclic nature of plant infection over decades [21]. This is believed to be caused by gene clusters on Phytophthora and plants, of Avr genes and their corresponding interaction partners in hosts, resistance genes [22]. Large shifts in plant genes where entire groups of coding resistance genes are gained or lost seem to be the primary culprit for both cyclic occurrences and widespread pathogenesis. This is a normal process in plant immunity where groups of up to 40 PAMP or similar gene groups are contained in cassettes. These are rearranged, translocated and lost or gained between successive generations [22,23,24]. Genomic changes affecting translation rates and alternative splice variants of proteins also play a role. Phytophthora then changes Avr gene sets, which seem to follow similar genetic mechanisms of clustering and rearrangement between generations. This has been termed the “genetics arms race” between host and pathogen for these specific pathogens [25,26]. Only if a particular resistance gene is present and properly expressed, will the host plant infer resistance [27].

Pathogenesis in half the cases has been shown to occur in pathogen associated molecular pattern receptors (PAMPs), often referred to as leucine rich repeat (LRR) proteins [28]. These same proteins inferring resistance also seem to be responsible in many cases for pathogenesis. Often only a small amino acid stretch, as little as 15, determines a plants resistance or susceptible through a respective Avr. Thus, only small interactions determine if pathogenesis occurs [29]. Widespread host target proteins rule out PAMPS as always being the resistance proteins involved, which are primarily the focus of most studies. However other proteins, such as E3 ubiquitin ligase, DAMPs and MAMPs are targets of some Avr [24,30,31]. These may be applicable to single Avr on gene clusters, and may be simply a more coordinated aspect of pathology as a whole, but have only been minimally researched [32]. A complete categorization or sub grouping for Avr proteins is deficient, due to lack of complete structural information and the difficulty in determination of plant host interacting proteins [26].

At the protein level, Avr were originally grouped based on N-terminal sequences termed RxLR motifs, and further sub-grouped from similar sequences at their C-terminal region in a few cases [25,33,34]. These allow large groupings, with putative similar functionality having as many as 60-70 proteins within a single group. Many of these are believed to be inactive at the transcription level, while functional proteins vary smaller sequence regions and single amino acid sites at a high rate [35]. Putative non-protein ligands include lipids such as inositol phosphate derivatives, varied ions, generalized membrane lipids and cholesterols [36]. Overall, the belief is that varied pathogenic proteins act in a coordinated way to modify one or several pathways in the host related to defense. Inhibiting or lessening responses significantly allows the pathogen to then thrive, reproduce and increase overall infection rates against a host organism. Dilemmas exist in reproducibility of ligand interactions, along with functionality of small regions of Avr proteins and knowledge of functionality. This lack of proper understanding of function, and conflicting data is the primary hindrance to furthering understanding at the protein mechanics level [37,38,39].

Here we focus on one protein from P.sojae termed avirulence protein 5 (PsAvh5) [40]. This protein has been shown to interact selectively with inositolphosphate-3’-phosphate (PI3P) lipids, however again this has met with reproducibility issues in basic kinetic experiments [41]. Additionally, it is believed this protein interacts with membranes of the host organism, and is secreted into the interhaustorial space by infecting Phytopthora. Experiments with related Avr3a, Avr3a4, Avr3a11 and Avr1b-1 from varied Phytopthora species have shown these proteins will accumulate in the host cell over time [42,43]. The highly homologous N-terminal region, containing a shared sequence motif of RxLR has also been shown to allow accumulation of fluorescent tags alone, pointing to a secretion or translocation mechanism. Despite this, the C-terminal region of PsAvh5 and the closely related proteins, without the motif, have been shown to confer susceptibility. This from both expression in host tissue in the absence of the pathogens, or applied from external application to tissue [40,44].

To address some of the unknown functionality, we employed a molecular structural approach. Phytopthora excreted proteins are problematic to study in vivo as they are secreted into interhaustorial spaces [45]. Haustoria structures are essentially embedded regions of the pathogen inside the host cell, making isolation of proteins involved in the infection processes difficult. In addition, these proteins have been shown to behave abnormally, cycling between soluble and membrane forms, and additionally between dimeric or non-dimeric states [46,47]. Both NMR and X-ray structures exist for closely related C-terminal domains of this subset of related proteins [40]. These allow a significantly advantageous starting point for our research objectives. We furthered these starting structures with completely refolded forms from prior computational experiments (Unpublished).

Starting with an acetylated refolded protein a series of Docking and Molecular Dynamics (MD) simulations were conducted. We initially test membrane interactions, and PI3P interactions using Docking. Additionally, it has been shown that in two studies these proteins may form dimers [25,46]. This has been controversial, as an equal number of publications indicate this may not be the case [37]. Using finalized monomeric models and protein-protein docking, we also explore dimerization in the same context. Through extended MD simulations, we also test the effects of models on different membrane types, primarily lipid rafts and general membrane architectures. This is a key area of focus, as little is known of the mechanisms underlying Avr attachment or entry into host cells at the amino acid level [33,48]. Using steered molecular dynamics (sMD), we also attempt to gain insight into kinetic processes involved with membrane lipids or ligands, to determine not only feasibility of models but interactions themselves [49,50,51]. Together, these data are utilized to shed light on several controversial and unknown functional aspects of PsAvh5, and to determine relevant interactions at the amino acid level necessary for the pathogenic process.

## Materials and Methods

All simulations were carried out using Gromacs 4.6.6 and 5.0.1, with a nose-hoover thermostat and parrinello-rahmen pressure coupling at 290 K and 1 bar respectively [52]. All equilibration runs used a V-rescale thermostat until consistent temperature was achieved, and then switched to a nose-hoover thermostat both at 290K. Models were all parameterized using a gromos 54a7 force field and SPC water [53,54]. Analysis of all data was conducted using Gromacs 5.0.6. An initial model of PsAvh5 acetylated at residue R 27, Ace-RTADTDIVYEPKVHNPGKKQVFIEDKLQKALTDPKKNKKLYARWYNSGFTVKQVEGGLDQ NENRELELTYKNLALGYAKYYQARRSQEAK, was used from prior refolding work (unpublished)

Initial dimerization was tested using Gromacs with different systems. In each a solvent system consisting of H2O, 0.1 M NaCl, 0.03 M KCl, 0.02 M MgCl, 0.01 M CaCl and 7 Zn ions, was used. In model 1, two monomeric proteins were placed in a solvent only unit cell 90 90 90, 4 Å apart. This was allowed to run for 600 ns, after an initial 50 ns equilibration. Two monomer proteins were also set on a membrane consisting of 300 1,2-dipalmitoyl-sn-glycero3-phosphocholine (DPPC) lipids and 100 cholesterol, in 90 90 120 unit cell, in the same solvent and allowed to run unrestrained for 1 us. Secondary dimerization was tested using the same monomeric PsAvh5 model docked to itself using the program HEX protein-protein docking with electrostatics and shape for several independent runs [55]. From the best-fit HEX docked model, the same solvent alone and solvent-membrane system consisting of DPPC were used to test stability. These were both allowed to run unrestrained for 200 ns each, after 50 ns equilibration runs. Another set of simulations for the HEX dimer and two monomers were then set up with 250 DPPC and 30%, 20%, 10% and 5% molar cholesterol. These were solvated using the same solvent, and allowed to run for 200 ns each, after a 50 ns equilibration run, to further test dimerization stability and interactions with membranes.

A more complex membrane set was generated to test membrane effects of both monomer and dimer models with dimensions 90 90 Å^2^. These consisted of 214 DPPC, 30 2-aminoethoxy-2,3-bis-octadec-9-enoyloxy-propoxy-phosphinic acid (DOPE), 40 or 15 Cholesterol, 7 Ceramide (18:1/24:0) and 5 2,3-bis(alkanoxy)propyl-2,3,4,6-tetrahydroxy-5-(phosphonatooxy)cyclohexyl-3-phosphate (PI3P) lipids. Lipids were parameterized to the 54a7 force field using the automated topology builder, and modifying files to match lipid atom names [56]. Membranes were allowed to reach stability over 100 ns unrestrained simulation time, and solvent of H2O, 0.1M NaCl, 0.03M KCl, and 0.02M MgCl. Membranes were assembled using overlaying sheets in gromacs, and g_membed [57]. All phosphorylated forms of the Inositol-phosphate head group were also tested through Docking, to the monomer and dimer structures. These included Monophosphate at the 2’,3’,4’,5’, 6’ position, bisphosphate at positions 2’-3’, 2’-4’ 3’-4’, 2’-5’ and 3’-5’, 2’-6’, 3’-6’, triphosphate 2’-3’-4’, 3’-4’-5’, 3’-4’-6’ and 2’-3’-5’ positions.

For both membranes the dimer, 2 monomers or no protein were generated and placed in solvent consisting of H2O, 0.1 M NaCl, 0.03 M KCl, 0.02 M MgCl, 0.01 M CaCl and 7 Zn ions, for a final unit cell of 90 90 120 Å^3^. Either 1 monomer or the dimer were first docked to the Inositol-3-phosphate head group, using both Autodock and Vina, then the model fit onto an already embedded PI3P lipid [58,59]. Structures were first analyzed with Pymol and Poisson-Boltzmann surface maps [60,61]. For placement of the second monomer, the protein was placed equidistant from the fit protein using half the box length along the X-Y membrane surface. Each membrane system was allowed to equilibrate for 100 ns, unrestrained after temperature and pressure were reached, with semi-isotropic pressure coupling. Final simulations were allowed to run an additional 50-100 ns unrestrained, and the last 30 ns used for comparative analysis using Gromacs internal analysis tools. Membrane density maps or thickness maps were generated using three dimensional 0.01 nm grid spacing, or single phosphate head groups between membrane leaflets respectively according to established methodology [62]. Where possible, maps were aligned using protein structures between successive frames at the bound PI3P molecule. Additionally, a visual attempt was made to initially align structures and all trajectory frames then fit using progressive fitting. All data and statistics were analyzed and plotted using Qtiplot© and Gnumeric©.

For steered molecular dynamics simulations, 6 evenly spaced starting models were taken from the last 20 ns of each unrestrained membrane simulation for the dimer or monomeric system [49]. This generated 12 starting models. Simulations were equilibrated for 4 ns, were set to reach maximum force at 15-18 ns, and reach a maximum distance of 2 nm from the membrane based on test runs with varied lipid restraints and pull forces (Sup.Movie 1, 2). These resulted in a simulation set with a harmonic pull force from 0 to 2000 KJ/mol/nm^2^ for 20 ns for covariance analysis. Force was applied to each protein in both states separately at the center of mass, providing two pull groups for each simulation. Lipids DOPE, DPPC and PI3P were restrained with a 1000 KJ/mol force in the Z direction initially at the terminal CH3 group, and simulations repeated using 2000 KJ/mol on the O1, O2 lipid position. Several types of trajectory analysis were conducted using Gromacs command line analysis tools, or other internal structural and simulation analysis tools according to published covariance protocols [50,63].

The software DSSP was used in conjunction with Gromacs and VMD for tertiary structure analysis across trajectories, and structures visually analyzed using Pymol and VMD [64]. Hydrogen bonding, bond lifetime, angle and distance were analyzed with Gromacs and cross-referenced with both VMD auxiliary tools and Pymol. Statistical analysis was also conducted using Gnumeric© and Qtiplot© from energies obtained through individual simulations, extracted through Gromacs. Total free energy (ΔG) for the PI3P, and Membrane and total change was also calculated using simulation times from 2-4 ns subtracted from the last 2 ns for each pulled simulation, using a running average system and means calculated in Qtiplot©, and checked against Gromacs calculations. Map matrix averages for both thickness and density were produced using auxiliary python scripts, Gnumeric©, and Gromacs. Plotting of data for surface maps was performed using Qtiplot©. Principle component analysis was conducted according to published protocols [63]. This included extracting energy vectors using Gromacs command line tools, in this instance the top 3 vectors corresponded to the X, Y and Z direction. These were then plotted and visualized using Qtiplot©. Images of trajectories were produced using Pymol and VMD from varied simulation single frames, or single trajectories for movies [64,65]. Surface electrostatics were analyzed using the advanced Poisson-Boltzmann software, and visualized using VMD [61,66].

## Results

### Dimerization does not occur with molecules in solvent simulations with submicrosecond simulation times

A starting monomeric protein, with N-terminal acetylation at the R27 residue, was duplicated and placed in random orientations 3-4 Å apart (Fig. 1A). These were embedded in solvent consisting of normal ion concentrations and allowed to run unrestrained for 200 to 300 ns. Despite several differing orientations, no complete dimeric protein became fixed. Proteins showed a slight propensity to align the α-helices from T95 to K116 in an anti-parallel fashion. These however never reached a stable state, with residues along the helices from either molecule moving 2-3 turns up or down the helix, with respect to the symmetric molecule.

**Figure 1:**
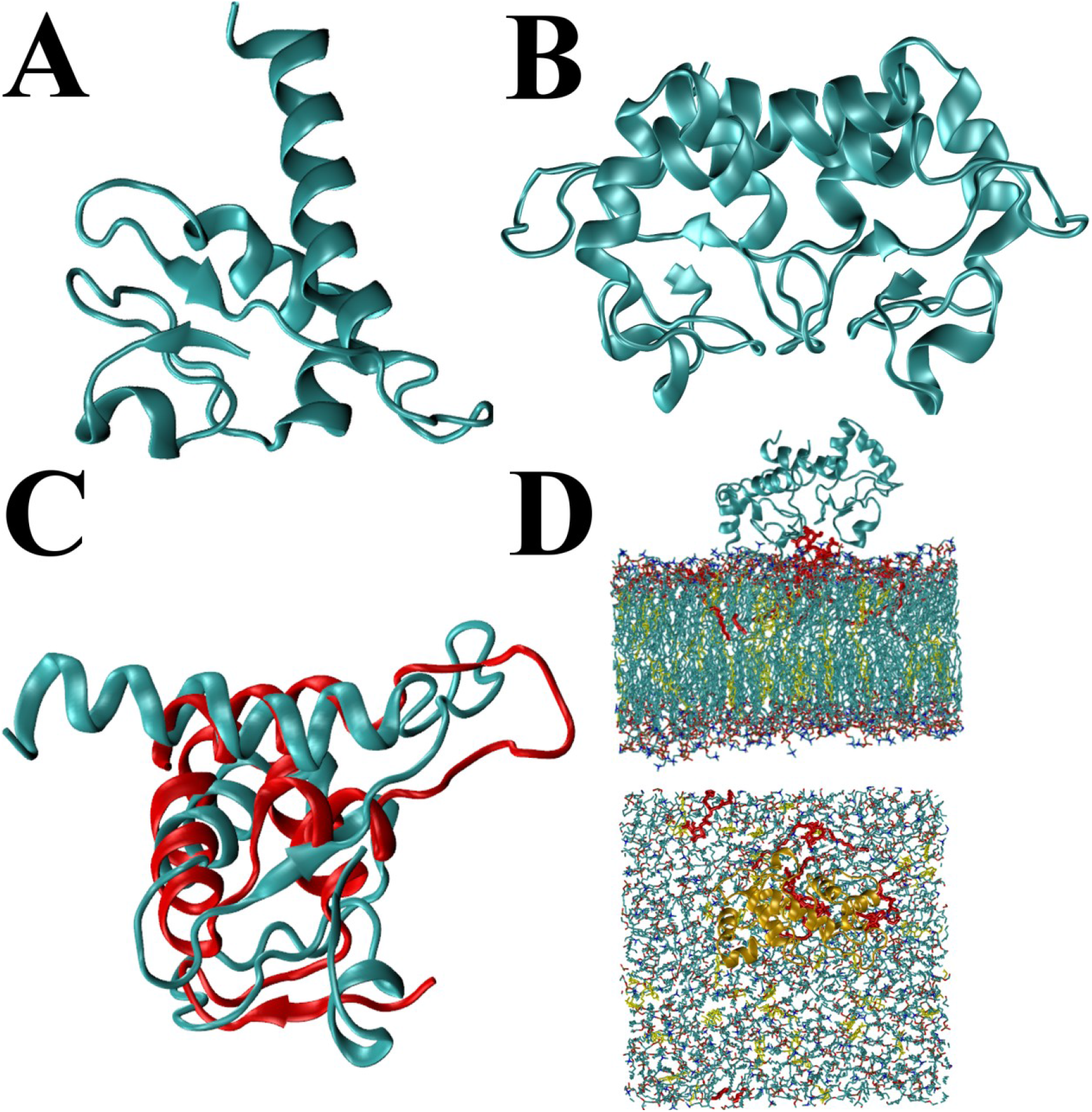
Starting Models. A) Final monomeric form of Avh5 obtained from solvent based refolding. N-terminus with acetylated Arg lower portion showing the 6 amino acid helix not previously determined in x-ray structures. Top, C-terminal Lys 116. B) Dimeric form of Avh5, lower portion oriented towards membrane. Visible ß-sheets are from Arg 27 and Thr 28 to Gln 79, Val 80 and Glu 81. C) Comparison of Avh5 (Aqua) and Avr3a11 from PDB 3ZGK (Red), showing the difference in the N-terminal residues. D) Starting models for Membrane simulations, with dimer shown. PI3P (red) DPPC, DOPE and Ceramide (aqua), Cholesterol (yellow) and Avh5 top (aqua), Bottom (Gold).

Using a membrane of 300 1,2-dipalmitoyl-sn-glycero3-phosphocholine (DPPC) and 100 cholesterol, simulations were continued for another 200 ns. This was repeated for 30%, 20%, 10% and 5% cholesterol for another 200 ns each. Two monomers were placed side by side with the α-helices from T95 to K116 arranged anti-parallel. This placed the region from R27 to H40 parallel to the membrane. Proteins slowly drifted apart on 10% and 5% cholesterol, and the α-helix in either protein from residue T31 to E36 became slightly oriented towards the membrane, with an approximate 20° angle offset, from residue D32 towards the membrane, and E36 slightly raised. A loop starting from A29 to T31 became embedded in the membrane surface, and T31 unwound shortening the helix by 1 residue. Hydrophobic residues F75, T76 and V77 also oriented into the membrane. Despite minor adhesion to the membrane, the two proteins did not achieve a stable dimer. Proteins drifted away from, and towards each other, but did not move further than 5-6Å apart. This was due to random longer charge based interactions between a large number of lysine, arginine and glutamic acid residues. It was found membrane simulations with 20% and 30% cholesterol had little lipid mobility, which also affected protein mobility. On these membranes, proteins did not move except for amino acid functional groups.

### Protein-protein docking utilizing electrostatics produces only one solution with high probability

To further test the possibility of dimerization, the software HEX was utilized using a combined electrostatic and shape based docking approach(107). This resulted in a single model, with lower scoring models only offset from the highest by 1-3 angstroms. Any larger changes in the output conformation resulted in 1000 KJ higher energies, although energies in docking algorithms are protein specific and comparative. In all docking runs, the same highest scoring structure was obtained even when docking parameters were varied. This highest scoring model was equilibrated in solvent for 20 ns, placed on a lipid bilayer consisting of 300 DPPC and 100 cholesterol in solvent, and then used for small molecule docking. The equilibrated dimer was able to remain folded, and showed slight differences between the two protein subunits, both between each other and between the monomer. Comparisons with the monomer after extended membrane simulations showed structural similarity to the related Avr3a11 (Fig. 1C). Simulations in solvent alone caused the dimer to begin to dissociate, and slightly move towards the monomer structure after 30 to 40 ns.

### Small molecule docking shows a single site on the monomer, and dimer that differ significantly

Autodock Vina was used to dock the PI3P charged head group alone, and PI3P with 3 carbon chains in several docking runs. The monomer and dimmer were used for separate runs, as the structures differed (Fig. 1 A,B). A single site not accessible in the dimer was found for the monomer with only a slight affinity above randomized docking. A mean value of 24.54 KJ/mol +/- 2.30 for the site, with background cut off 20 KJ/mol for randomized affinity. In the dimeric form, a novel site is formed between the two dimers, from residues N72 to K78, with H40 and Q79 also participating. This dimeric ligand site showed a strong affinity of 42.66 KJ/mol+/- 6.70 for IP3P only. The site itself can orient a single PI3P in two equally affine conformations, rotated by 180°. Affinities were taken from the Autodock internal scoring function, and are based on point electrostatics, idealized gas states and desolvation using generalized tables. As a result, affinities are only comparative, and used for screening of small molecules. Additionally hydrogen bonding of less than 4 hydrogen bonds was used for screening cutoff. No other derivatives of inositolphosphate were able to dock into the dimer structure. All derivatives showed an affinity for the monomer in the same site, however were just below the 20 KJ/mol cut off.

### Structural differences occur between the dimer subunits, and between PI3P bound or unbound monomers

To test the feasibility of each model obtained, a more complex set of membranes were utilized. These contained either 5.5% or 13.3% molar ratio of cholesterol. This was used to distinguish affinities for lipid rafts or general cell membranes. Each membrane also included 213 DPPC, 30 2-aminoethoxy-2,3-bis-octadec-9-enoyloxy-propoxy-phosphinic acid (DOPE), 7 Ceramide (18:1/24:0) to maintain surface charge and 5 2,3-bis(alkanoxy)propyl-2,3,4,6-tetrahydroxy-5-(phosphonatooxy)cyclohexyl-3-phosphate (PI3P) lipids, with 15 or 40 cholesterol respective of molar ratio. Each membrane initially started with dimensions of 92×92 Å^2^ with a Z axis of 145 Å, and was solvated. After equilibration, each unit cell had shrunk to 83.74 x 86.59 x 150 Å or 86.56 x 89.50 x 142.60 Å for the 5.5% and 13.3 % cholesterol respectively.

Using the best docked conformation for the PI3P head group, the monomer and dimer were then fit onto the membrane by aligning the head group to an embedded PI3P. It was found after 80 ns of unrestrained equilibration for membranes alone, PI3P molecules would cluster in groups of 2-3 around Ceramide lipids. This caused a clustering of PI3P prior to addition of the proteins. To maintain comparison, the same PI3P on either membrane was used for the bound monomer and dimer (Fig. 1D, Sup.Fig. 1). In all further analysis, these were used to center trajectories as closely as possible. An additional, unbound monomer was also added to the membrane from a randomized orientation obtained in the DPPC only simulations.

Each membrane simulation was allowed to equilibrate for 70-120 ns, unrestrained for each of 6 membrane systems. These constituted membranes alone, membrane with PI3P bound dimer or membrane with bound IP3P monomer and unbound monomer simultaneously. At the end of equilibration, the software DSSP with Gromacs, and VMD for cross reference, were used to analyze tertiary structure for the entire protein in each simulation from the terminal 30 ns. Differences between monomers were primarily in order of secondary loops, and the β-bridges or sheet from residues R27 through A29 to Q79 through E81. For the dimer, PI3P makes interactions with both molecules simultaneously. The overall dimer structure has several hydrogen bonds with molecule A, and only 2 hydrogen bonds with the mirrored protein. Our initial structure placed both molecules in a position superposable by 180° around a central axis in the Z direction from the membrane. After equilibration, the molecules differ slightly, with an axis from molecule A to B along the X-Y plain at 30° from the membrane. Intermolecular β-bridges between symmetric V77 and Q79 are also no longer present in the PI3P bound form. This is primarily caused by Molecule A slightly embedding T31 through I33, and G43 through F48 into the membrane, the later as an entire loop. The symmetric molecule only makes transient interactions with residues in these loops and the membrane. Additionally, molecule A changes the helix from residues T31 to E36 into a 3-10 helix. Additionally, F75 becomes embedded into the membrane, while held in a tight β-sheet within the adjacent molecule B.

### Monomeric and dimeric proteins have significantly different effects on membrane structure

To test effects on membranes directly from protein, each simulation was continued for an additional 50-100 ns after each equilibration run. Thickness and density maps were generated for both cholesterol types from monomers, dimers and a control membrane without protein. Average thickness for dimers was close to that found in the control sets, with slight differences in PI3P only (Fig. 2). Overall control and dimer containing membranes have a mean thickness of 39.5 and 36.8 Å +/- 1.1. Dimers tended to raise PI3P to the mean level or slightly higher by 1.5 Å in 5.5% cholesterol membranes only, for the bound PI3P. Monomers uniformly dropped the entire membranes average thickness by 1.1 Å and 1.4 Å respective of the 5.5% and 13.3% cholesterol membranes from control membranes. Additionally, the entire membranes were more uniform across the entire plain.

**Figure 2:**
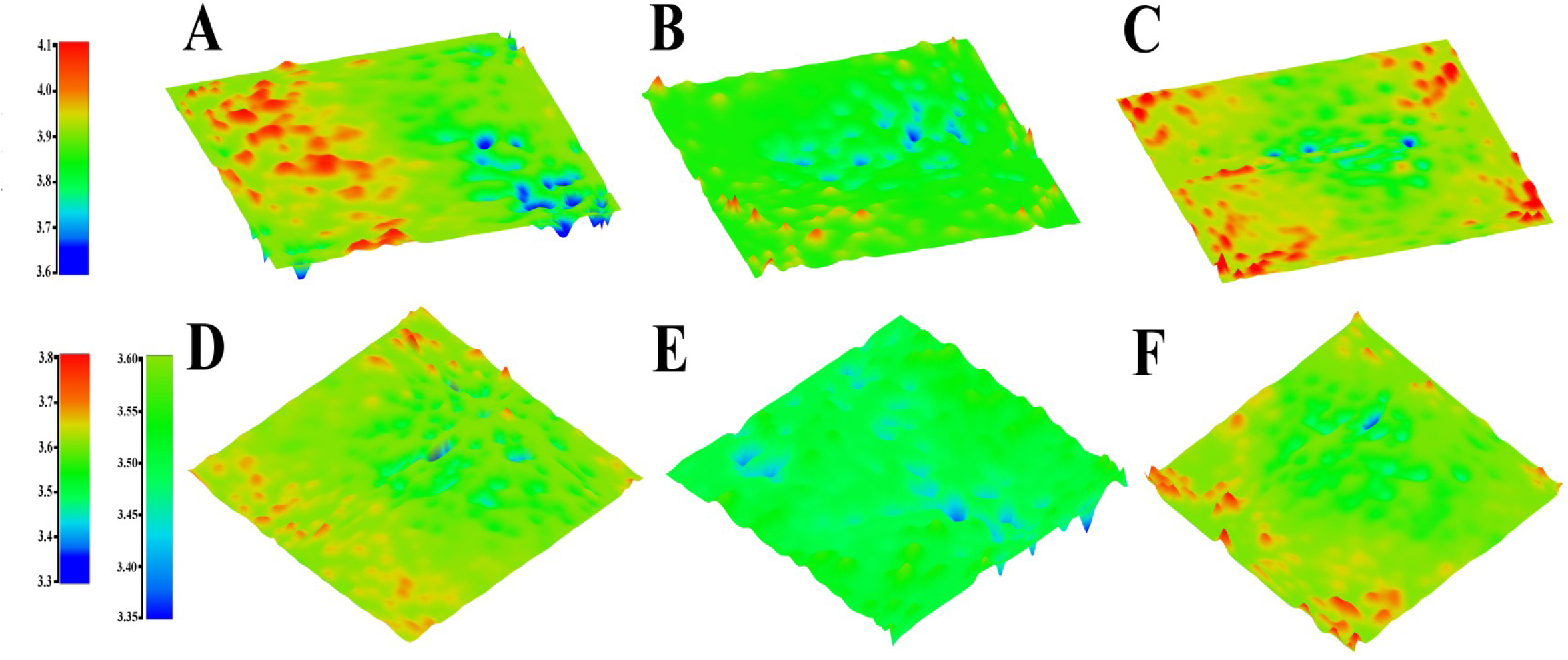
Thickness Maps of Model Membranes from Extended Simulations. Top, 5.5 % Cholesterol membranes, Bottom, 13.3 % Cholesterol Membranes. Color bars to left are in nanometers. A) Control membrane, B) Monomeric Avh5 C) Dimeric AVh5, D) Control membrane, E) Monomeric AVh5, F) Dimeric Avh5. The second color bar to the left of D-F is an expansion of the first color bars, as E) shows more subtle differences do to a uniform membrane. In both sets, the monomeric Avh5 shows a larger effect on the membranes than the dimer. For B, C, E, F protein is aligned to lower thickness areas. In B, E, bound PI3P is thinnest (blue) peak. For C bound PI3P is higher (yellow, orange) peak next to lowest peak (blue) and D, mean thickness (no obvious peak). In C, F lowest (blue) peak beneath low hydrogen bonded Avh5 subunit.

In the control or dimer membranes, a thickness was determined by clustering of the PI3P lipids to single Ceramides that caused clustering which appear as blue pits. In the dimer, the Ceramides migrated away from the PI3P via repulsive charges from the protein, however the clustering of PI3P remained due to interactions with the protein. These were transient with non-bound PI3P, lasting shorter than 1 to 2 ns, however longer range electrostatics along with these cyclic interactions from lysines maintained them near the protein. With the monomer, PI3P clustering was disrupted and lacked longer hydrogen bond formation with PI3P. For PI3P and other lipids, the monomer orients a series of alternated charged residues, Tyr, Lys, Arg, Glu and Gln towards the membrane. Single hydrogen bonds are formed, and the alternating charge acts to pass lipids to the next residue allowing the protein to glide over the surface of the membrane. This in effect caused a uniform thickness to occur across the membrane by homogenization of lipid-lipid contacts, and an almost uniform thickness forced underneath the protein.

With the dimer, membrane interactions were a combination of hydrophobic and charged residues, Several DPPC lipids were slightly raised 2 Å to 4 Å, through interactions with residues forming two pockets that lasted 4 to 8 ns. The two longest lived pockets were formed by residues F75, K44 and G43 which were able to fix DPPC-cholesterol pairs for 8 ns, and F48, V47 and K45 which fixed single DPPC or DOPE lipids for 4 to 6 ns. This effectively utilized lipids as hydrogen bonded extensions of the dimer, fixing it to the membrane, with much less movement of lipids within van der walls cut off of the protein than seen with the monomer.

Density maps were generated for the entire membrane constituent atoms, and aligned to the thickness maps (Fig. 3). For the 5.5% cholesterol membranes the control, monomer and dimer mean were 13×10^-23^, 16×10^-23^ and 14×10^-23^ g/nm3. The 13.3% cholesterol showed mean densities of 13.5×10^-23^, 17×10^-23^ and 15.5×10^-23^ g/nm3 for the control, monomer and dimer. The monomer containing membranes showed the highest degree of density variation, almost fluctuating for each lipid pair across the membrane sinusoidally. In contrast to the dimer, the monomer containing membrane showed no higher density peak correlation with cholesterol coordinates (Sup.Fig. 2). For the control and dimer membranes, higher peaks correlated to cholesterol coordinates in both membrane sets. Structurally the difference can be explained by pinching of the membrane by the monomer, but not the dimer structures. This effectively bunches the lipid tails adding space between adjacent tails for each lipid, yet forcing many to adopt non-linear shapes (Sup.Fig 3). Membrane pinching was mirrored only slightly in the dimer membrane, only showing a density variation of roughly 2×10^-23^ g/nm3 while maintaining the overall membrane thickness observed in the control membranes. Visually, the differences are noticeable with the dimer and control membranes having a much more linear lipid tail placement perpendicular to the membrane plain. Overall, the difference in dimer containing membranes densities was slight and unexpected, while the mean higher difference from the monomer containing membranes correlates well with the thinner thickness observed as each membrane contains the same number of lipids and cholesterols in each simulation.

**Figure 3:**
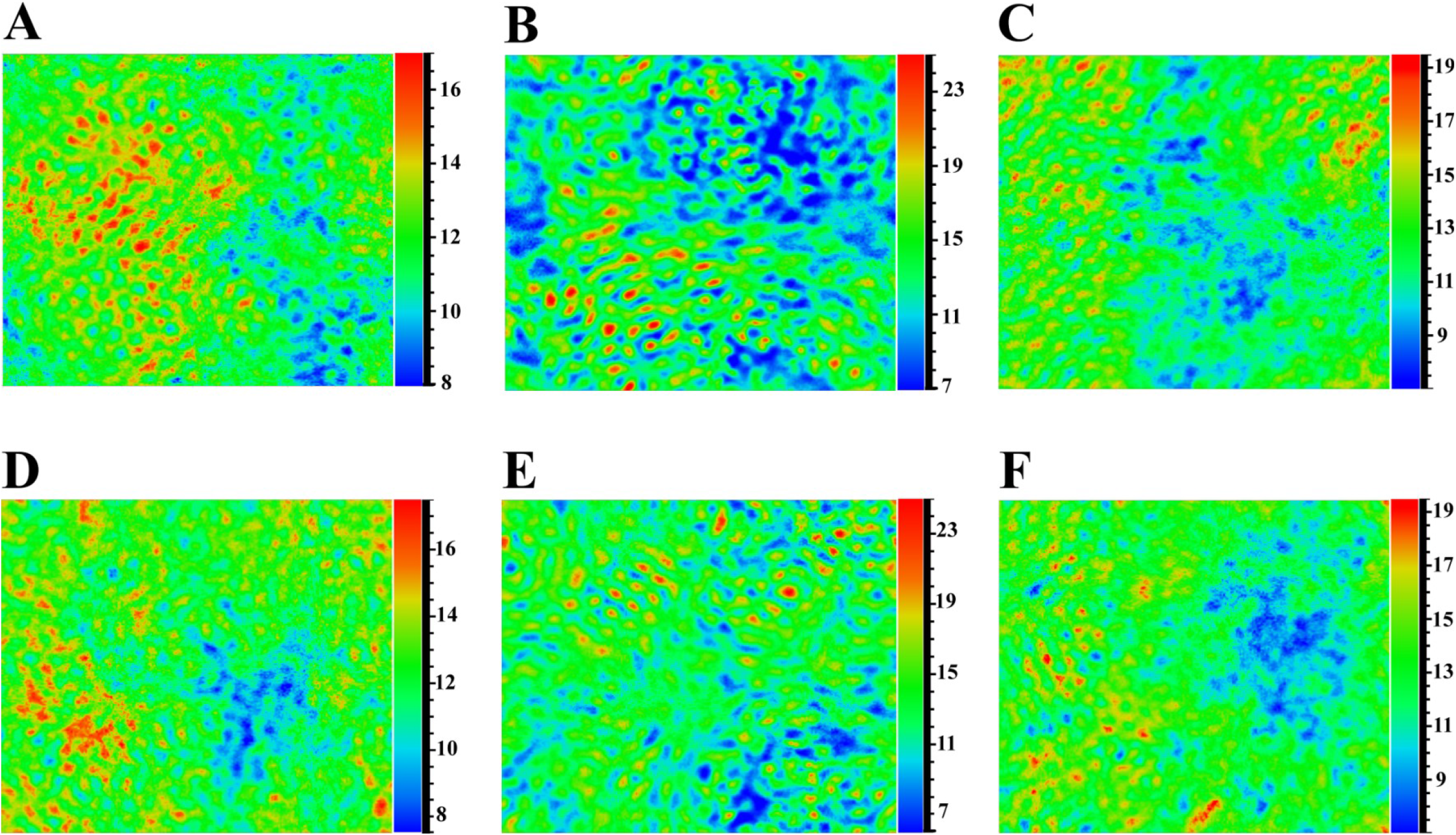
Density Maps of Model Membranes from Extended Simulations. Top, 5.5% Cholesterol membranes, Bottom, 13.3% Cholesterol membranes. Color bars to right of each graph are specific to each plot. A) Control membrane, B) Monomeric Avh5, sitting on upper blue portion, C) Dimeric Avh5, also aligning to blue portion. D) Control membrane, E) Monomeric Avh5, bound Avh5 on blue portion, second unbound on green adjacent, F) Dimeric Avh5, aligning to blue region. In both controls, the PI3P clusters to the blue regions and 3 Ceramide molecules on the top surface. For monomeric Avh5, a single PI3P forms the darkest blue peak, while the remainder are evenly spread. For Dimeric Avh5, the bound PI3P is held at the mean, and the remainder cluster around the Avh5 dimer.

### Lipid Hydrogen bonding differs significantly between the monomer and dimer proteins

Hydrogen bonding was analyzed for membrane and PI3P specific interactions across unrestrained extended simulations (Sup.Table 1). Differences occurred between all residues involved, in addition to length of interactions for each protein. For the monomer, the site determined from docking is not exposed to the membrane except through interactions from the main chain backbone atoms (Fig. 4A). In total 10 hydrogen bonds between 1.8 Å and 3 Å occur across the simulation. Main hydrogen bonding occurs between the functional groups of R27 and the O of the 3-phosphate, K52 and 5-O position of the Inositol, E81, G82 and Y35 with O atoms in the lipid head portion of the PI3P. The remainder of the hydrogen bonds form and break cyclically over the simulation, ranging from 2-8 ns in duration. Amino acids Q86 and K38 hydrogen bond with the inositolphosphate portion of the PI3P, rotating slightly to accommodate either residue during simulations. These do not occur simultaneously, but alternate every 2-8 ns.

**Figure 4:**
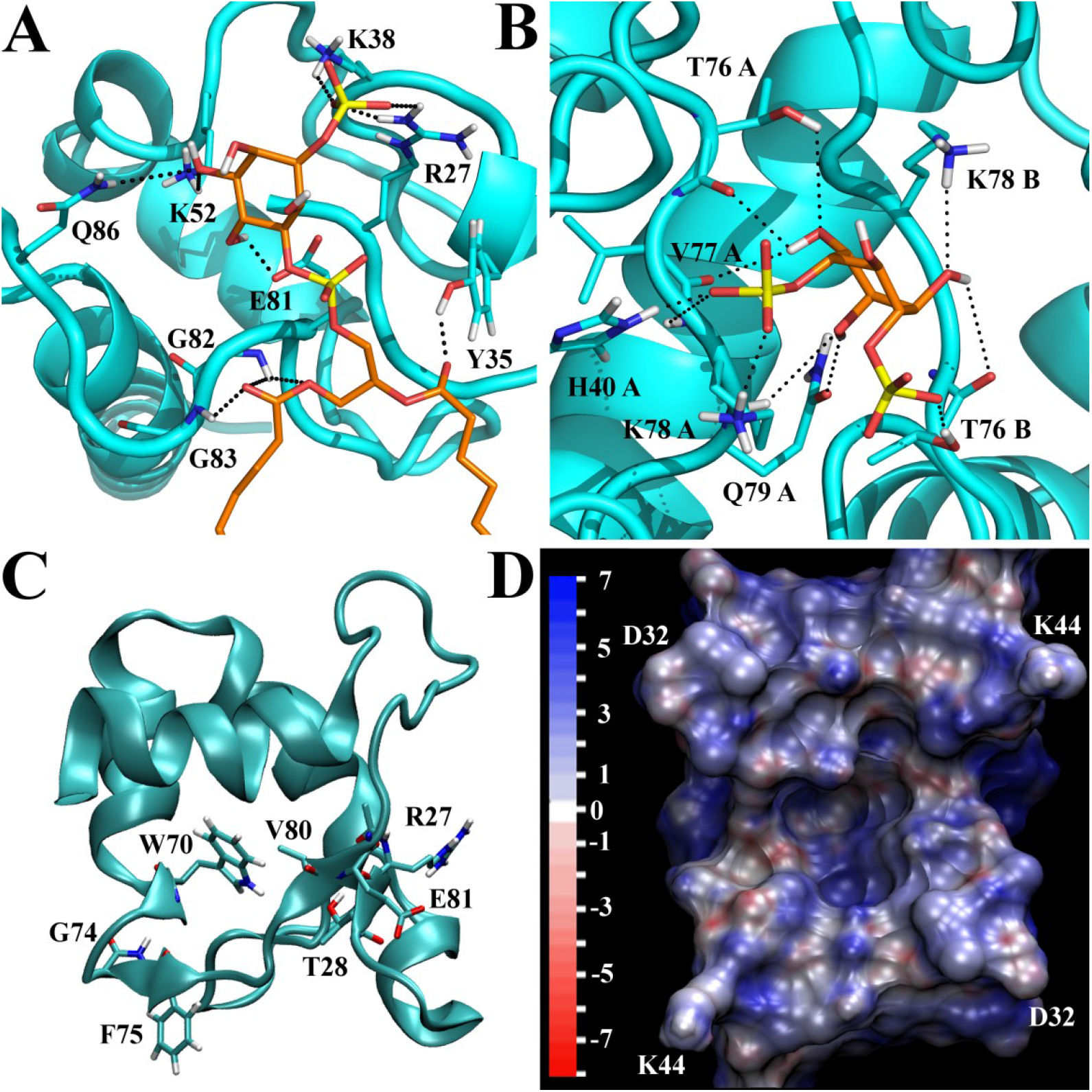
Functional Sites on Monomeric and Dimeric Avh5. A) Monomeric binding site found from docking, with bound PI3P. B) Dimeric binding site from docking with bound PI3P. C) Sliced view of subunit from Dimeric Avh5, showing small ß-sheets at either end of stretch from Phe 75 to Gln 79, bottom oriented towards membrane. Model shows Molecule B subunit with less hydrogen bonds to PI3P. D) Electrostatic surface of dimer, scale in eV. Model is from bottom facing membrane towards the viewer. The upper portions of the dimer were sliced away to show binding pocket (center). Model is from bound dimer after equilibration on membrane. In A, B, hydrogen bonds shown are composite from last 30 ns of extended simulation (not pulled), and do not all occur simultaneously.

Dimer hydrogen bond formation with PI3P occurs in a pocket formed from dimerization (Fig. 4B). Structurally, the binding site found in the monomer is partially inaccessible to any lipids, including the PI3P, being buried within the protein-protein interface (Fig. 4C). Effectively, the structure is analogous to a clamp when IP3P is bound. Upon extended unrestrained simulations, the PI3P forms the majority of hydrogen bonds with one molecule, and occupation excludes the 180° mirrored site from being occupied. This is illustrated in a charge surface map, which shows the binding pocket formed (Fig. 4D), that can orient the Inositol group rotated 180° with equal affinity, but has a phosphate site specific to a single protein in the dimer when bound. Structurally, this site can only sterically allow PI3P, or IP3,6P, the latter of which does not occur often in nature nor dock into the site. However, once bound the second phosphate site is lost through slight structural changes.

Hydrogen bonds are formed with the backbone of T76, V77 and K78 in the bound protein, with the mirrored V77 being forced upward in protein B. Specific interactions occur with H40 and K78 functional groups and the O groups attached to the 3-phosphate. Additionally, T76, K78 and Q79 functional groups interact with the OH and O groups of the Inositol, maintaining the entire PI3P in a rigid conformation fixed into the dimer pocket. The residue T76 also pulls the phosphate on the lipid portion into an orientation slightly higher than the mean lipid phosphates in the membrane, and remains fixed for the majority of simulation time. An adjacent K78, from protein B, forms transient bonds with Inositol O groups, along with the backbone atoms from T76 in the same protein. This K78 residue also probes the membrane adjacent to the PI3P, with bonds between the Inositol oxygens lasting 3 to 6 ns, and random membrane orientations for 3 to 4 ns.

### Dimeric protein-protein hydrogen bonding greatly affects protein subunit structure

Dimerization of the protein allows the helix region between residues K52 through T58 to extend by three residues, to I49. The terminus of this region also places F48 and V47 directly against the membrane. With bound PI3P, the occupied molecule A embeds the loop between G43 to F48 into the membrane, well below the amide groups of the lipids. This loop acts as a key anchor region, sequestering two lipids and fixing a cholesterol molecule as an anchor, directly hydrogen bonded to the protein backbone O and NH between F48 and V47 through the terminal OH. Terminal helices from T95 to K116 form a zipper arrangement of alternating charges, and several hydrophobic residues placed away from solvent between molecule A and B. An additional large patch of these helices have 8 tyrosines buried in the internal region of the protein directly below and away from solvent, and above loops S73 to Q79. These loops are stabilized by 2 asparagines and 2 glutamines in a symmetric hydrogen bond network between both molecules.

Notable differences occurring between dimer molecule A and B, and between IP3P bound monomer, were found in β-sheet or β-bridge arrangement. In molecule A, the structure is more rigid, with beta bridges between R27 and E81, and Q86 to L94. In the IP3P bound monomer alone, a shift to bridging E88 to L94 for the later bonding occurs, opening the loop to more solvent. Not in the monomer, a 3-10 helix also occurs from residues T31 to Y35 in molecule A. In molecule B, β-sheets replace the bridges. In the non-IP3P monomer simulations, only the first set of bridges becomes a sheet. Sheets occur with residues R27, T28, A29 to Q79, V80, E81, and between W70, Y71 to F75, T76. This later W70 precedes a α-helix and acts to hold the internal loop from F75 to G82 more ridged than the counter molecule. More stable hydrogen bonds that re-occur cyclically or lasting longer than 6 ns between molecules in the dimer are listed (Sup.Table 2)

### Differences in total energy changes upon membrane binding show dimeric PsAvh5 prefers lower cholesterol, while monomeric forms have a lipid raft preference

For monomeric forms of PsAvh5, PI3P was the major contribution to total energy change. This was determined for total interaction energies, and total free energy changes inclusive of solvent effects (Table 1). Comparatively, the monomeric form had roughly twice the energy associated with membrane interactions over the dimeric form, even though most interactions were non-specific single hydrogen bonds. Only a slight difference was observed between 5.5% and 13.3 % cholesterol, which can be attributed to hydrogen bond lifetime. These were greater in the higher cholesterol concentrations, however not significant enough to show a preference for membrane types in the IP3P bound form. The non-bound monomer has a slight preference for lipid rafts, which is observed primarily in the Z direction vectors moving away from the membrane.

**Table 1:** Total Free Energy Changes for Steered Simulations. Total free energy changes inclusive of solvation energy and protein self interactions calculated from steered molecular dynamics, and averaged from 3 separate simulations for each model. In each, Mol A bound PI3P or high hydrogen bonded subunit, and Mol B unbound PI3P or low hydrogen bonded subunit. First two, monomeric form, second two dimeric form. Free energy was calculated using PIP3 or Lipids inclusive of solvent effects for “PI3P “ and “Membrane” columns. ΔG_Total_, total Gibbs free energy inclusive of every atom in the protein and solvation of membrane and protein components. The dimer was treated as a single protein for Gibbs calculations. Energy in Kjoule (Kcal).

|  |  |  |  |
| --- | --- | --- | --- |
|  | **5.5% Cholesterol** |  |  |
|  | **IP3** | **Membrane (Lipids)** | **ΔGTotal** |
| **Mol A** | **-299.45 +/- 9.69 (-71.57)** | **-153.91 +/- 18.64 (-36.79)** | **-738.92 +/- 26.30 (-176.61)** |
| **Mol B** | **-51.92 +/- 16.07 (-12.41)** | **-31.02 +/- 20.82 (-7.41)** | **-491.39 +/- 17.40 (-117.45)** |
| **Mol A** | **-129.16 +/- 21.89 (-30.87)** | **-30.20 +/- 6.94 (-7.23)** |  |
| **Mol B** | **-27.58 +/- 8.94 (-6.59)** | **-5.10 +/- 1.08 (-1.22)** | **-918.54 +/- 106.62 (-219.54)** |
|  | **13.3% Cholesterol** |  |  |
|  | **IP3** | **Membrane (Lipids)** | **ΔGTotal** |
| **Mol A** | **-134.64 +/- 16.54 (-32.18)** | **-3.68 +/- 0.35 (-0.88)** | **-707.02 +/- 107.46 (-168.98)** |
| **Mol B** | **-74.43 +/- 9.04 (-17.79)** | **-77.45 +/- 26.68 (-18.51)** | **-648.80 +/- 44.35 (-155.10)** |
| **Mol A** | **-103.91 +/-5.27 (-24.84)** | **-21.07 +/- 1.86 (-5.04)** |  |
| **Mol B** | **-12.84 +/- 4.15 (-3.07)** | **-4.42 +/- 0.17 (-1.06)** | **-534.98 +/- 15.94 (-127.86)** |

A much greater effect was observed for the dimeric form. In regards to the bound PI3P alone, the higher bound molecule A roughly tripled its interaction energy with lower cholesterol levels. For the symmetric molecule B, the effect was largely from random lipid interactions within the membrane. Molecule B also showed higher energy with lower cholesterol levels. This is contradictory to what is expected from higher lipid mobility, and seems to be related to flexibility of the molecular pair on the surface of the membrane. As the second molecule becomes tilted by 20-30°, there is much more variation in the angle for the 5.5% cholesterol membrane surface interactions, and slightly greater distances by 1-2 Å. This in turn allows longer interaction with lipids than when held more rigid on lipid raft higher cholesterol levels, with movement of the dimer across the membrane.

Overall, when solvation is accounted for the monomeric PsAvh5 and the dimeric have relatively equal affinities for PI3P and membrane systems. The total free energy changes correlate with other findings, in regards to the dimer preference for lower cholesterol. Dimeric preference for membrane types are just below the significant difference level, with ANOVA values of P <0.04. Monomeric forms show no significant difference in affinity between membrane types if PI3P is included, but a strong preference for non-IP3P containing monomers for lipid rafts. In both energy comparisons, this is also reflective of the standard deviations obtained. Parallel simulations in solvent alone also show the dimer will slightly unfold the terminal helix, and residues F48 to D51. Additionally the two molecules become slightly rotated from each other, suggesting the dimeric form is only stable on membranes. A higher affinity found in the monomeric PsAvh5 for PI3P is partly due to orientation alone. The Inositol and 3-phosphate group overhangs the backbone of residues from V80 to G83, and steric hindrance rather than direct hydrogen bonding causes a larger specific affinity. Dissociation of the IP3P is from hydrophobic effects in the lipid tail, rather than hydrogen bonding to the protein. This indicates the molecule must undergo a temporary structural change to initially bind PI3P.

### Monomeric and dimer proteins show similar spatial mean displacement vector peaks corresponding to the same residues or regions

Surprisingly both the monomeric and dimeric forms of PsAvh5 show the highest spatial changes in similar residues (Fig. 5). These are in W70, the loops beginning with K44 and K45 in both molecules, also the loop V80 to L84 and Q86 to R90. These are associated with β-bridges or β-sheets at end residues, and for V80-L84 a hydrogen bond from the loop against a helix. These are disrupted when the molecules move into solution. In both K45 interacts with a backbone O on ASN 41, while K44 interacts with lipid molecules. This loop region containing K44/K45 is unstructured in monomeric forms absent of IP3P. The ß-bridges or sheets also show peaks for residues R27 through A29, and hydrogen bond with residues in the helix from K52 through K55 respectively. The only slight differences in vector peaks was found in the helix region from T31 to E36 for the dimeric protein, loops from F75 to Q79 from the PI3P binding site in the dimer, and D85 to L94 which forms a compact loop in the dimer. In higher cholesterol, the monomeric form with IP3P mirrors the dimeric changes from F75 to Q79 and D85 to L94. These correspond to the largest vector peaks in angstroms overall. Main differences observed translate into more compact loops in dimeric forms, which exist as membrane embedded loops in monomeric forms. Structural differences observed between regions from W70 to L94 highlighted in the vector map, show the monomeric form adopts a larger change across all amino acids moving into solvent.

**Figure 5:**
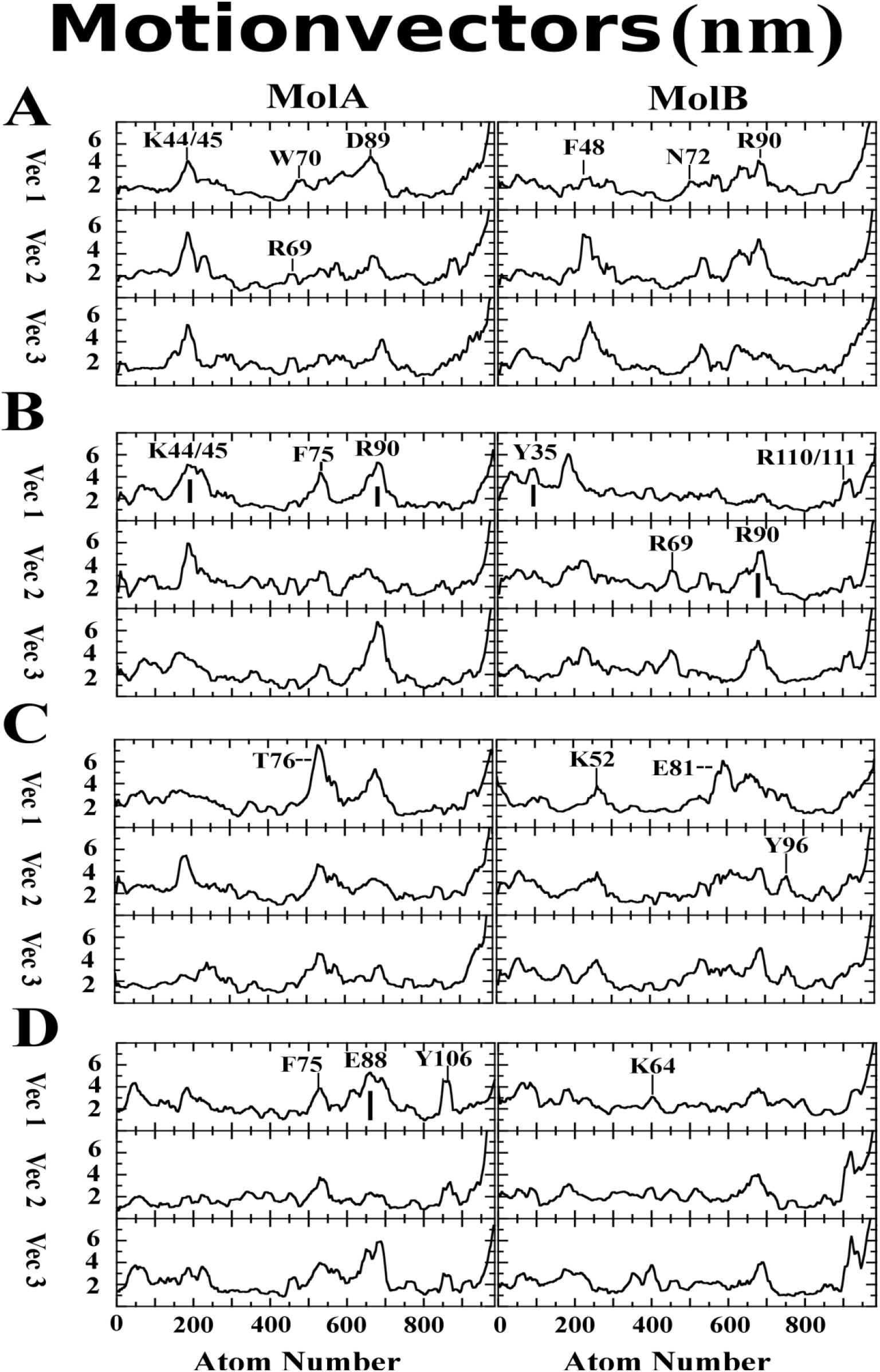
First Three Spatial Eigenvectors from Steered Molecular Dynamics Simulations for Avh5. Means from steered molecular dynamics simulations for first 3 highest spatial eigenvectors corresponding to the X, Y and Z directions over 20 ns simulations. A) Monomeric Avh5, Mol A bound PI3P and Mol B unbound. B) Dimeric Avh5, Mol A high hydrogen bound PI3P and Mol B low hydrogen bonding. A, B both on 5.5% cholesterol membrane. C) Monomeric and D) Dimeric Avh5 as in A and B, on 13.3% cholesterol membrane. Various Amino acid regions are labeled at highest motion peak, thin |, above, or thick | labeled below due to space, − −, label from side due to space in graph. A background of 2 nm has not been removed from random motions. Scales are in nanometers.

### Energy projections show larger differences in eigenvectors between dimer and monomeric PsAvh5 functional amino acids

Eigenvectors were filtered to remove background random fluctuation, and then projected with energy values associated with each motion. Correlated energy and motions were then analyzed using principle component (PC) analysis plots of the first three vectors, which account for 90% of the total energy change within the proteins (Fig. 6, Sup.Fig. 5). These were roughly equal in energy content of 30% +/- 1.50 total energy, signifying lack of larger structural changes. Total energies for individual vectors only varied by roughly 0.5 %, and were ranked computationally by energy order. Directional components for each vector correspond to all motion in the X, Y and Z plane for vector 1, 2 and 3 respectively. Less overlap was observed between monomer and dimer PsAvh5 than with the spatial eigenvectors alone.

**Figure 6:**
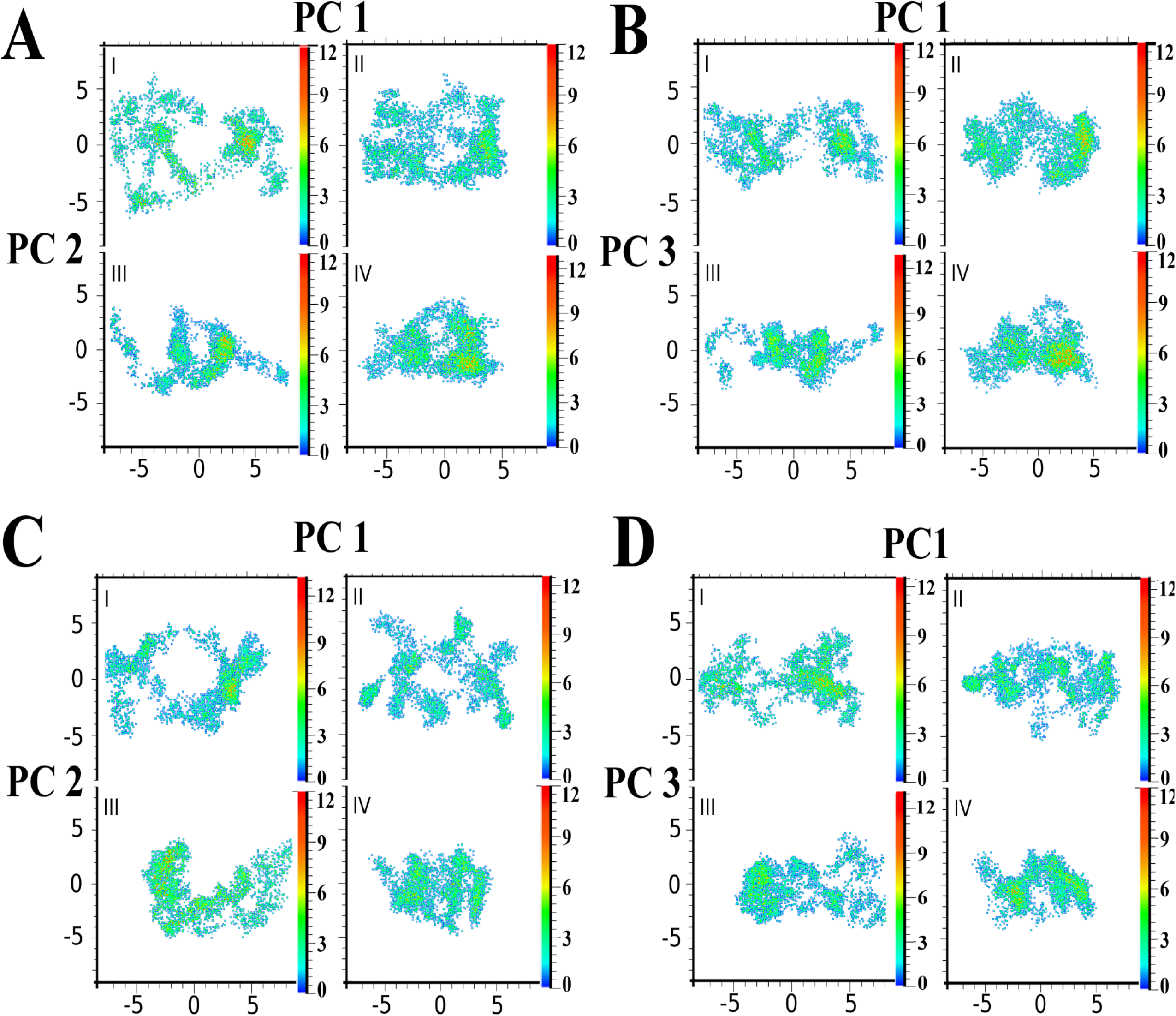
Energy Principle Components of Steered Molecular Dynamics Simulations. Total free energy changes, for the first three eigenvectors (90% total energy) for each Avh5 model. Eigenvectors 1-3 corresponding to PC1-3, were in the X, Y and Z directions respectively. All are on same scale X-Y in nanometers, with X-axis component labeled (top) and y-axis component labeled (right). Energy bars to right are scaled for individual PC plot, in Kcal/mol, as they vary slightly in total amplitude. I, Monomer IP3P bound, II, Dimer Molecule A, III, Monomer, no IP3P, IV, Dimer Molecule B. In the dimer, Molecule A is higher hydrogen bonded with IP3P, and Molecule B lower. A,B) 5.5% Cholesterol, C,D) 13.3% Cholesterol.

For dimeric PsAvh5 on 5.5% cholesterol membranes, the PI3P bound molecule showed the highest peaks for G74 through V77 that correspond to the re-arrangement of a β-bridge and rearrangement of the loop from F75 to Q79 to accommodate the PI3P. These are apparent around the 1 nm to −1 nm Y-axis and 3nm X-axis coordinates on PC1 versus PC2 plots, and again in the PC3 plot (Fig. 6 A-II and B-II). For the symmetric PsAvh5 subunit, two peaks were observed, with residues Q79 and symmetric V77 changing to a β-bridge, as well as the re-arrangement of the same loop molecules and the helix from T31 through E36 changing from an α-helix to a 3-10-helix. Coordinates are roughly scattered from 1 nm to 3 nm in the X-axis, by −1 nm to −3 nm Y-axis for the Helix. This later is present in molecule A, however does not show as high an energy peak, as residues from T31 to E36 already form a 3-10-helix arrangement. More distal lower peaks in molecule A correspond to lipid interactions, primarily with the loop region around K44 and K45. In molecule B, these appear as much more ordered interaction, and include contributions from W70 and Y71 with β-sheet to β-bridge rearrangement at −1 nm to −3 nm X-axis and 0 nm to −2 nm Y-axis (Fig. 6 A-IV).

For correlated PC1 and PC3, the helix from T31 to E36 shows a larger peak in molecule B than with the second vector, signifying for this molecule a greater force is found along the Z direction perpendicular to the membrane (Fig. 6 B-IV). This is also apparent in the loop regions from F75 to Q79 for both molecules, with molecule B showing much higher energy peaks in this region, as it forms a β-sheet from β-bridges. For molecule A, a third peak becomes apparent for R69 through N72, due to the loss of the β-sheet to a β-bridge arrangement between W70 and Y71 with G74 and F75, and the region containing loop K44 and K45 has a much more focused peak at −5 nm in the X axis, and 0 nm on the Y axis. Differences between PC2 and PC3 correspond to a shift towards more centralized individual amino acids interacting with the membrane, as well as the same residues involved in the PI3P binding process already identified. These occur only slightly above the already determined peaks, from 4 nm to 5 nm X-axis and 0 nm to 3 nm Y-axis coordinates. Overall, molecule A shows a more compact set of interactions due to a more rigid structural arrangement around IP3P, and a large peak associated with the maintained β-bridge from W70. Coordinates are close to the PC1 and PC2 plot.

Cholesterol effects are much more apparent from PC plots than other energy methods. On the 13.3 % cholesterol membranes, both subunits on the dimer shift the majority of energy components away from the helix region between T30 and E36, which are much broader on plots spatially. Additional lower energy contributions are seen for molecule A from the loop between G43 and I49, with a shift to more specific amino acid or amino acid pairs. These appear as more focused isolated cluster regions in PC1 and PC2 plots (Fig. 6 C-II). Focused peak regions run the edge area occupied by the loop between F75 to Q79, and include I49, L45 and H40 (Fig. 6 C-II). Symmetric PsAvh5 has no peaks for these regions, showing a majority of energy still associated with the helical region T30 to E36, the same stretched areas from F75 to Q79 and β-sheets as observed with the 5.5% cholesterol membrane. For plots of PC1 and PC3, the energies are roughly the same as with the 5.5% cholesterol for both subunits, however become much more residue specific with molecule A. Both molecules show noticeable lower energy. In molecule B, there is also a large loss of energies for the loop region between F75 through Q79 (Fig. 6 D-II, VI). While these residues still contribute, the orientation of this molecule in the PI3P bound dimer is much more rigid, making membrane interactions less. Overall weaker hydrogen bond formation even when present, and a slightly more rigid molecule that does not change residue conformations as drastically, yield a weaker but superimposable map between different membranes based on cholesterol (Fig. 6 B, D).

In monomeric forms of PsAvh5, energy correlations peak only for residues between F75 and Q79. No interactions occur with PI3P and these residues, however the stretch is directly placed on the membrane surface. Adjacent stretch of residues from V80 to D85 is also placed against the membrane, while remaining above the helix T30 through E36. A remainder of peaks are scattered single amino acids or amino acid pairs adjacent to each other in the PI3P bound monomer. These correspond to hydrogen bonds found in simple analysis across trajectories (Sup. Table 1). Unbound monomer matches molecule B from the dimer. Differences are observed from 0 nm to −5 nm Y-axis and 1 nm to −3 nm X-axis. Here, the higher peak set corresponds to amino acids V80 to Q86, while still including the region T30 to E36. (Fig. 6 B-II).

Effects with increased cholesterol are a much more focused peak set for individual amino acids or amino acid pairs in higher concentrations. Observable differences between energy distributions are a slightly more compacted protein and a preference in total energy for the unbound molecule towards higher cholesterol. This later is apparent only when the Z directional component is involved in plots with PC3. Additionally, the IP3P unbound monomer shifts energy contributions to the α-helix T95 to L116 from residues placed against the membrane. These are observed between - 1 nm to −3 nm X-axis and −1 nm to 3 nm Y-axis (Fig. 6 C-III).

Peak overlap between monomeric PsAvh5 and dimeric forms corresponds to only minor conserved structural similarity in subunit architecture. A comparison with DSSP tertiary structures in the membrane bound, IP3P bound forms highlight the areas found in energy peaks upon dissociation with the membrane (Fig. 7). Comparatively, PC analysis shows the dimer has more energy attributed to the Z vector direction, perpendicular to the membrane. Conversely, monomers have most energetics applied parallel to the membrane. This is more apparent in comparisons of 5.5% cholesterol maps than higher cholesterol concentrations. Higher peak areas do not overlap between monomeric and dimeric forms. Most structural regions are in completely different environments between monomer and dimer, being in direct contact with lipids in one environment, and buried internal or against solvent in the other. Still, some homologous overlap occurs, primarily with the loop F75 to Q79. Overall, the dimeric form acts more mechanical, while the monomeric form shows energies primarily with direct lipid interaction.

**Figure 7:**
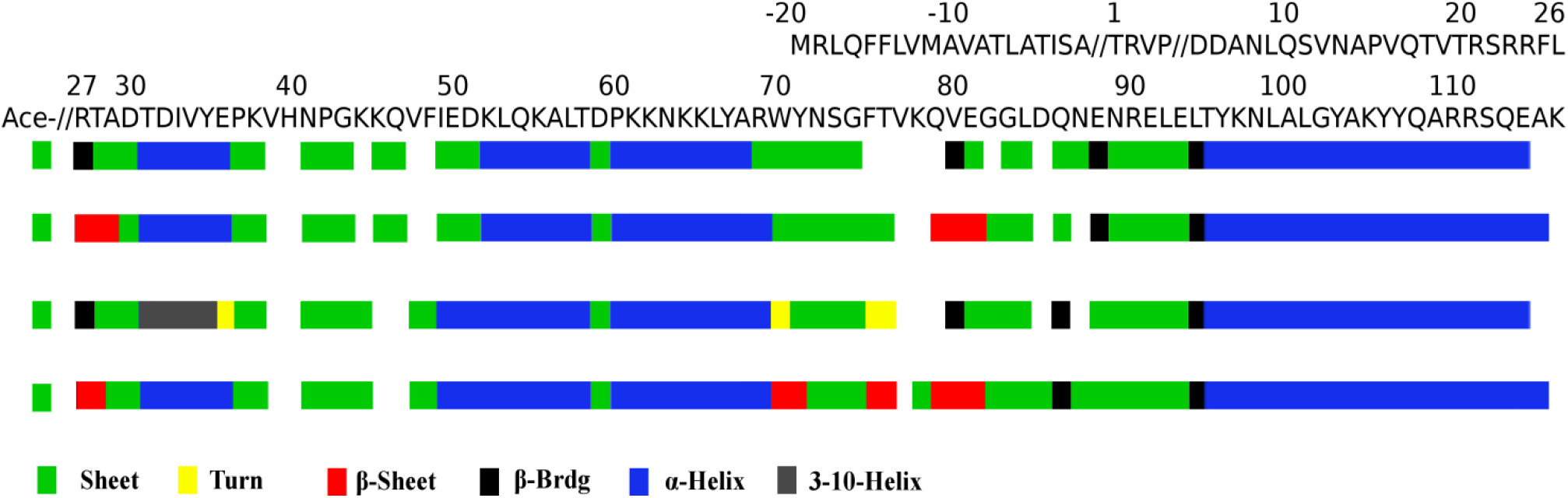
Pali (DSSP) Tertiary Structure Map of Avh5 Forms. Sequence of Avh5 shown at top corresponds to color code in legend. From top to bottom, Avh5 monomer with bound PI3P, Avh5 with no bound PI3P, dimer Molecule A subunit with higher hydrogen bonding to PI3P, dimer Molecule B subunit with lower PI3P hydrogen bonding. The initial sequence starting from Thr 1 is also shown at top, //, cleavage site. Ace, Acetyl group, shown as sheet and separated from Arg 27.

## Discussion

Understanding Avr proteins and their role in plant pathology is of key interest to several economic sectors, and includes national defense [10,67]. Understanding these proteins at the molecular level can provide the foundation needed to accomplish desired goals of all these areas. Using structural features, from dimeric or protein interactions with plant resistance proteins, as well as smaller molecular ligands has already been highlighted as a targeted area [25,68]. These features narrow the necessary research needed to elucidate cellular mechanisms lying between the molecular level and organismal based research now being conducted. Additionally, structural information allows more rapid rational design approaches to be utilized. Thus reducing computational requirements by providing focused targeted areas on proteins to combat food and crop losses [69,70]. These single studies can be expanded to similar molecular sets within the pathogen, when key structural and functional comparisons are utilized from the wealth of already available genomic data.

Taken together, competing models emerge from our simulation studies. Both models have affinities for the membrane equal to other known membrane surface binding molecules of similar size [71,72]. In the first model, the PsAvh5 only exists as a dimer, and this dimer interacts with PI3P. Initially dimers would be drawn to PI3P rich regions of a membrane, leaving when the ligand is found into general membrane regions. In a slightly more complex model, PsAvh5 exists as a monomer in solution, forming a dimer in the presence of PI3P after transiting the interhaustorial space. A final third model is even more complex. In this model, the PsAvh5 exists initially as a monomer secreted into the interhaustorial space of invading Phytophthora. As a monomer, the PsAvh5 binds to and releases membranes until it finds the proper region, along with a second PsAvh5 protein. Multiple PsAvh5 would congregate causing slight invagination of host cell membranes, possibly close or on lipid rafts. Eventually, the PsAvh5 re-orients two subunits into a functional dimer. In the dimeric state, the PsAvh5 then initially finds a PI3P and its respective PAMP or host protein ligand. From some studies, there is evidence that Avr may have different functionality within the same protein at different cycles of the infection process [36,73]

Functionality suggests the dimeric form of the protein more closely reflects a realistic PsAvh5 model. Partially this is based on the binding site formed, as it is completely shaped to the Inostiol- 3-phosphate head group, and would hinder other forms of inositol groups (Fig. 4 B, D). Additionally, there is a functional mechanic process initiated by the binding of PI3P, suggestive of a necessary liganded state found in other studies. Also, it has been shown in other studies PsAvh5 binds solely to IP3, and H40, which bends to clamp the IP3P in place in our structure is important [36,40,44,74]. In solution, both molecules of the dimer hold the IP3P binding site open, primarily through a 3-10-helix. When bound, one of these re-arranges into an α-helix further clamping onto the head group of the ligand, and forming β-sheets creating a more rigid subunit. Our data shows this IP3P binding process to be mechanical, suggesting it is not a random artifact from modeling alone.

Inclusive of all data analyzed, the process follows the following mechanism for dimers. The ligand PI3P is initially recognized by T31, K78 and K45 and moved to the central portion between anti-parallel loops from F75 to Q79. This is accomplished by bending of the helix between T31 and E36 from a fixed 20° angle against the membrane to a more direct placement, with T31 oriented towards the membrane and E36 away. In this molecule, the helix also changes to an α-helix, and together these orient the H40 into a position to form hydrogen bonds with the 3’-phosphate of IP3P. Aiding in this process the β-sheets between residues R27, T28, W70, Y71, G74 F75, Q79, V80, and E81 change to single β-bridges. This in turn allows the two loops to form a more stretched orientation in the more affine PsAvh5 subunit, from F75 to Q79. A net effect is to orient the top surface of the dimer, consisting of 4 α-helices, at a 30° angle from the membrane, and clamp onto IP3P as a membrane anchor. Analogous residues on superimposed Avr1b (Fig. 1 C) or Avr3a4 have shown some of these to be functionally relevant [34].

Monomeric PsAvh5 models are more viable in solution, however have an equally strong affinity for membranes. Stability suggests these may still be a realistic form in some functional aspect. The lack of specificity in the PI3P docking site indicates this site may be an artifact, or may be used as a generalized binding mechanism to any phospholipids with an adequate head group. This is also apparent from the hydrogen bonds formed, with many being to lipid oxygens, found in a wide range of lipids (Fig. 4 A). The monomeric form of PsAvh5 is stable in a more compact form throughout extended solvent only simulations. In total, these were around 1.2 us in solvent systems, and indicate it is a stable structure. In contrast, the dimeric PsAvh5 will begin to unravel without a membrane substrate in 40 ns, and dissociate the two subunits. Structural simulations show an unstable protein will not last for a few hundred nanoseconds, and are used as tests for properly folded proteins [75,76,77]. Additionally, the non-IP3P bound form has a slight preference for lipid rafts, which has been shown in some publications. Like most of the research in the field, this again has met with disputed reproducibility or conflicting results.

In differing models, the missing factors are the known effector target proteins if any, excretion mechanisms from pathogen and location for the formation of dimeric structures. For PsAvh5 and structurally related homologous these are PAMPs in some cases, but other host proteins have been implicated as targets [40,78](31,130). Protein have been shown to accumulate in host cells, however a number of conflicting results exist. These proteins have also been shown to accumulate in pathogen Golgi, and very little has been studied on proteins in houstorial space. In part, this is due to the difficulty in studying native proteins in these structures, as they are surrounded by host tissue. Most studies have relied on tags, co-expression of molecules or expression within host or cells alone. Thus, a definitive model of the protein throughout the ineffective process from Phytophthora is difficult to conclude. Conclusive of our data, both structures are feasible as far as stability is concerned but the dimeric form fits much more of the published data for PsAvh5 regarding biochemistry. For both the monomer and dimer forms, the affinities found are equal to comparable membrane surface binding proteins ().

Future directions can be more focused based on the findings presented here. The data provides a more feasible approach to explain or fashion mutational studies to determine processes involved in parthenogenesis. Additionally, specific sites can be explored chemically, through massively parallel docking of compound libraries. These can form starting points for further pesticide development, or novel control methods. It has been shown through similar studies, docking can provide molecule libraries and these in turn used to test computationally determined sites through inhibition assays. Compounds can be tested for function at both cellular levels, as well as through simple in vitro kinetic assays. Structural information can also be utilized to create models for related Avr proteins. Overall, our data adds to research indicating PI3P is a ligand, and that PsAvh5 or related proteins form dimers interacting directly with membranes.

## Acknowledgements

We would like to thank the following for allowing the work to be conducted between 2014 and 2015. Pennsylvania State University, ICDS-ACI, Computations for this research were performed on the Pennsylvania State University’s Institute for Computational and Data Sciences’ Roar supercomputer, and Botany and Plant Pathology Dept. Oregon State University Super Computing Cluster, Corvallis, Oregon, and Oregon State University, Dept. of Botany High end computer and, the Oregon State University College of Agricultural Sciences, Dept. of Botany and Plant Pathology, staff for allowing the research to be conducted.

---

**Supplemental Figure 1:**
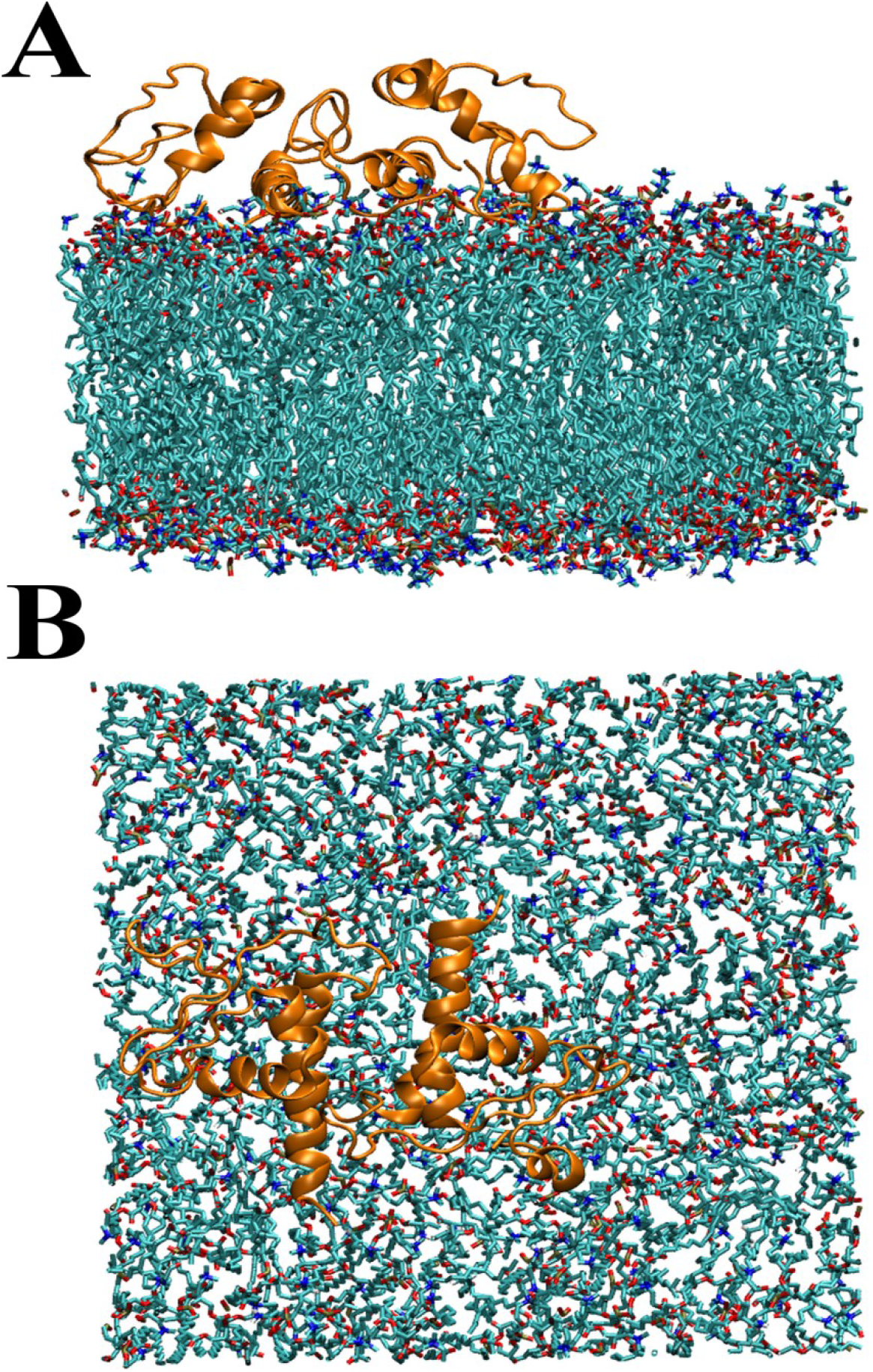
Starting Steered Membrane Simulations for Monomeric Avh5. Shown is Avh5 with bound PI3P (left) and unbound (right) at beginning of steered molecular dynamics simulations. After extensive unrestrained runs, monomers will aggregate. Avh5 (gold) on top of membrane lipids (aqua). Head groups are colored by atom type, oxygen (red), nitrogen (blue) and phosphate (yellow).

**Supplemental Figure 2:**
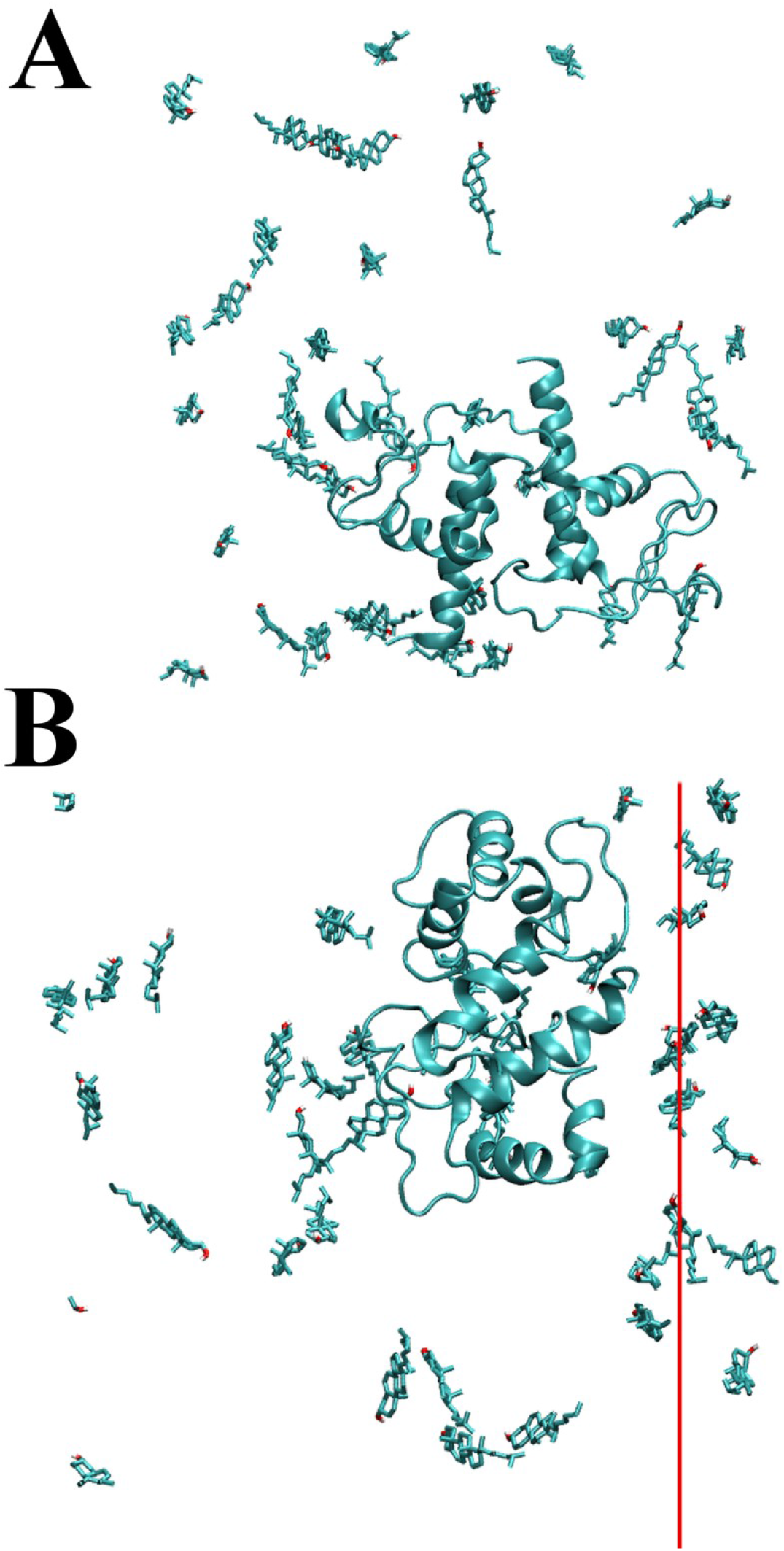
Cholesterol Positions in Extended Simulations. A) monomeric and B) dimeric Avh5 models shown from above. In B, red line shows periodic boundary change, with all atoms right moved to left side of box for maps in Figure. In A, Avh5 with bound PI3P at right. And in B, lower Avh5 subunit high hydrogen bound. Both are 13.3% cholesterol membranes.

**Supplemental Figure 3:**
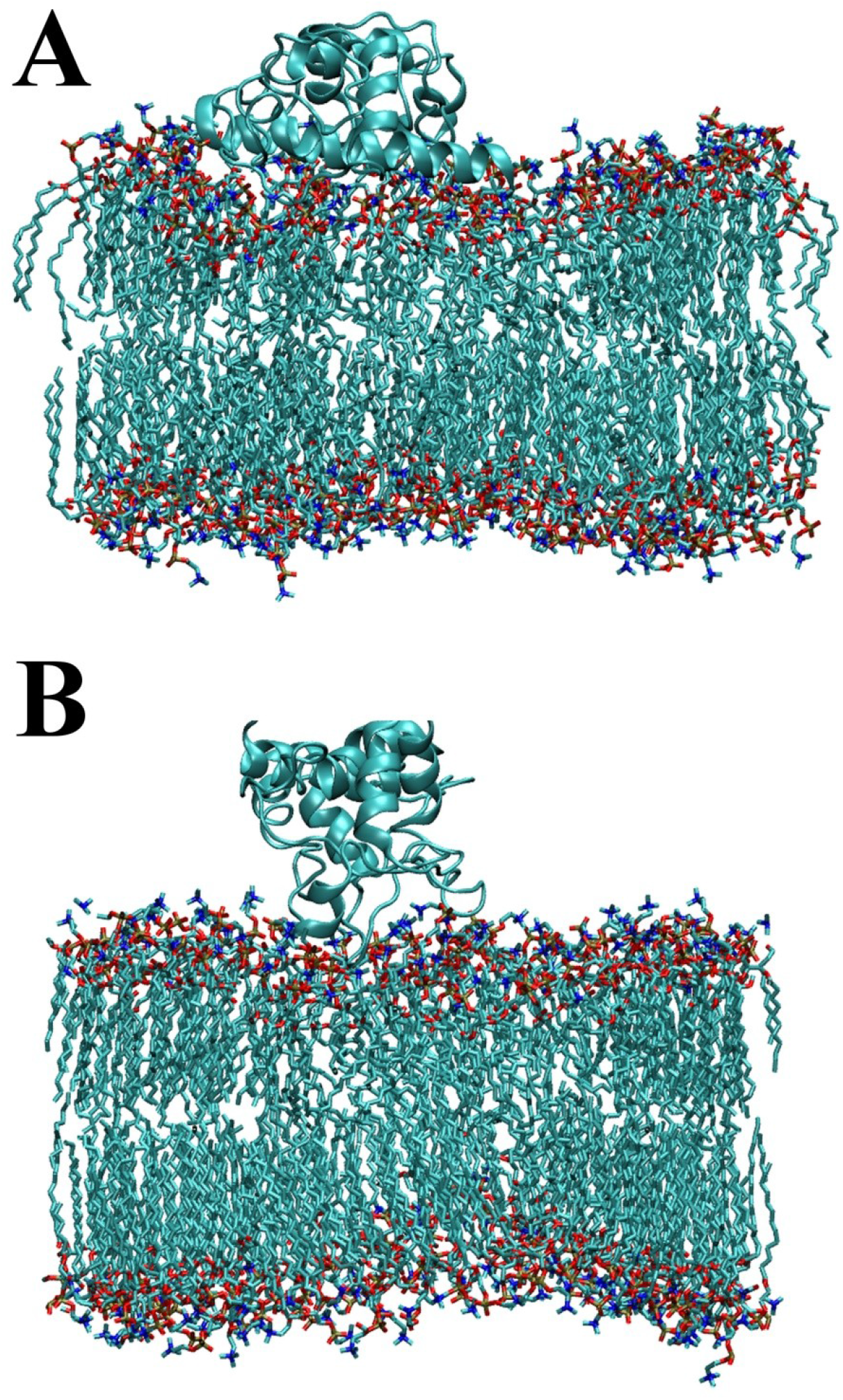
Lipid Tail Order in Extended Unrestrained Simulations. Single frame snapshots from end of 50ns unrestrained simulations for A) monomeric and B) Dimeric Avh5. In A, PI3P bound monomer away from viewer, and B, PI3P bound subunit left. In monomeric forms, the lipid tails adopt more randomized orientations, while in dimeric forms tails are more aligned for extended stretches. Both are 13.3% cholesterol membranes.

**Supplemental Figure 4:**
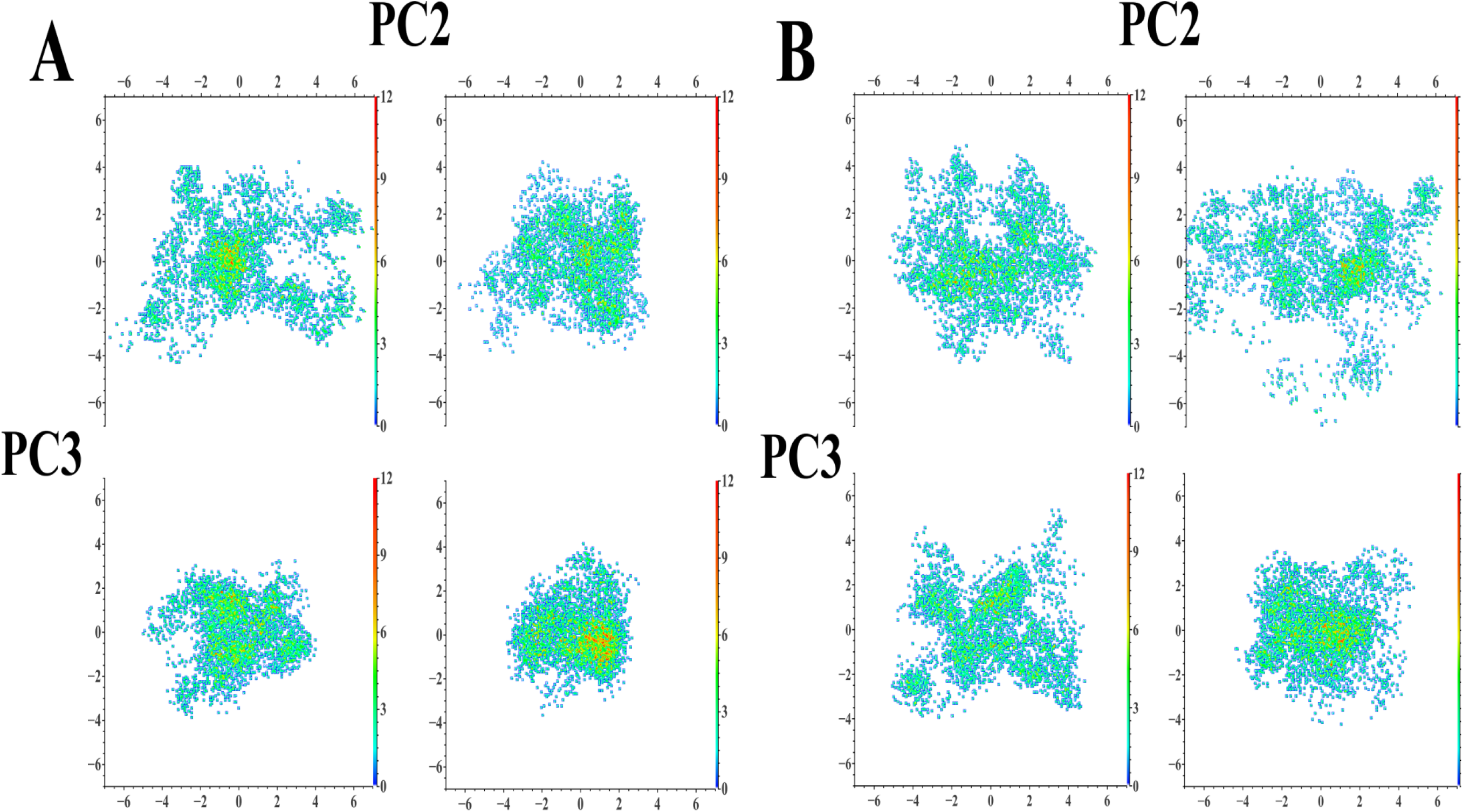
Principle Component Plots of Eigenvectors 2 and 3 for All Avh5 Models. Shown as in Figure 8, principle component plots for vectors 2 (top) and vectors 3 (left) for all models. A, 5.5% cholesterol membranes and B) 13.3% cholesterol membranes. Top, PI3P bound monomer or high hydrogen bonded subunit, bottom, unbound or low hydrogen bonding subunit Avh5. Left, monomeric and right, dimeric forms of Avh5. Heat map bar to right of each set for individual graphs shown. Vector lengths are in nanometers for X, Y-axis

**Supplemental Table 1:** Protein-Membrane Hydrogen Bonds Across 50ns. All hydrogen bonds formed during 50 ns of unrestrained simulation on respective membrane type. Monomeric with PI3P bound (Mol A) and unbound (Mol B), and dimeric high hydrogen bonded to PI3P (Mol A) or low hydrogen bonded (Mol B). “Number of Different Lipids” column indicates number of individual lipids bound during entire simulation, and “Lipid Type” list of types bound during simulation. The lipid PI3P is listed only for transient interactions, as 4 unbound PI3P are also in each membrane. None of the unbound PI3P became bound in the monomer, or due to steric hindrance in the dimer. All PI3P interactions behaved like other lipids, as single hydrogen bonds. First sets, left are for 5.5% cholesterol membranes, right, 13.3% cholesterol membranes for both monomer and dimer. Total, total hydrogen bonds formed during simulation with lipids.

| Monomers | Mol A residues | Lipid Type | Number of Different | Mol B Residues | Lipid Type | Number of Different |
| --- | --- | --- | --- | --- | --- | --- |
|  | (Bound IP3P) |  | Lipids (50ns) | (No IP3P Bound) |  | Lipids (50ns) |
|  | Thr 31 | DPPC | 1 | Thr 31 | DPPC | 2 |
|  | Asp 32 * | DOPE | 1 | Asp 32 * | DPPC | 1 |
|  | Ile 33 * | DOPE | 1 | Val 34 * | DPPC | 1 |
|  | Glu 36 * | DOPE | 1 | Tyr 35 * | DPPC | 1 |
|  | Tyr 71 | DPPC | 1 | Tyr 71 | DOPE/IP3P | 2 |
|  | Asn 72 | DPPC/DOPE | 3 | Ser 73 | DPPC | 2 |
|  | Ser 73 | DPPC/DOPE | 4 | Thr 76 | DPPC/DOPE | 2 |
|  | Thr 76 | DPPC | 2 | Lys 78 | DPPC/DOPE | 3 |
|  | Lys 78 | DPPC | 2 | Gln 79 | DPPC | 1 |
|  | Gln 79 | DPPC | 1 | Gly 83 | DPPC | 1 |
|  | Leu 84 | DPPC | 2 | Asn 89 | DPPC | 1 |
|  | Asp 85 | DPPC | 1 | Lys 97 | DPPC/IP3P | 5 |
|  | Tyr 96 | IP3P | 2 | Tyr 103 | IP3P | 1 |
|  | Lys 97 | IP3P | 1 | Ala 104 | IP3P | 1 |
|  | Tyr 103 | DPPC/DOPE | 2 | Tyr 107 | DOPE | 1 |
|  | Lys 105 | DPPC/IP3P/Ceramide | 3 | Gln 108 | DPPC/IP3P | 2 |
|  | Gln 108 | DPPC/DOPE/IP3P/Ceramide | 4 | Arg 111 | DPPC/DOPE/IP3P | 6 |
|  | Arg 111 | DPPC/DOPE | 4 |  |  |  |
|  | Ser 112 | DPPC | 1 |  |  |  |
|  | Gln 113 | DPPC | 1 |  |  |  |
|  | Ala 115 | DPPC | 1 |  |  |  |
|  | Lys 116 | DPPC | 1 |  |  |  |
|  |  | Total 40 |  |  | Total 33 |  |

Supplemental Table 1: Protein-Membrane Hydrogen Bonds Across 50ns
| Dimers | Mol A residues | Lipid Type | Number of Different | Mol B Residues | Lipid Type | Number of Different |
| --- | --- | --- | --- | --- | --- | --- |
|  | (Bound IP3P) |  | Lipids (50ns) | (No IP3P Bound) |  | Lipids (50ns) |
|  | Thr 31 | DPPC | 2 | Asp 30 | DPPC | 1 |
|  | Gly 43 * | DPPC | 1 | Thr 31 * | DPPC/IP3P | 2 |
|  | Lys 44 * | DPPC/Cholesterol | 5 | Asp 32 * | DPPC/IP3P | 2 |
|  | Lys 45 * | DPPC/Cholesterol | 3 | Ile 33 * | DPPC | 1 |
|  | Gln 46 * | DPPC/DOPE/Cholesterol | 3 | Val 34 * | DPPC/IP3P | 2 |
|  | Val 47 * | DPPC/Cholesterol | 2 | His 40 | DPPC/IP3P | 2 |
|  | Phe 48 * | DPPC | 1 | Lys 44 | DPPC | 1 |
|  | Ile 49 * | DPPC | 1 | Lys 45 | DPPC | 1 |
|  | Lys 52 | DOPE | 1 | Ser 73 | DPPC | 1 |
|  | Gly 74 | DPPC | 1 | Lys 78 | DPPC/IP3P | 3 |
|  | Phe 75 * | DPPC | 1 | Gln 79 | DPPC/IP3P | 2 |
|  | Arg 90 | DPPC | 1 |  |  |  |
|  |  | Total 22 |  |  | Total 18 |  |

**Supplemental Table 2:** Dimer Intermolecular Hydrogen Bonds. List of all strong hydrogen bonds between subunits in dimer, lasting for majority of 50 ns unrestrained simulation on membranes. Both cholesterol membrane types showed same hydrogen bonding. In table, Mol A, high hydrogen bonded to PI3P subunit, and Mol B low hydrogen bonded. “Non-symmetric”, strong hydrogen bond formed during course of simulation not present in symmetric side of dimer. “ß-Bridges Formed Without IP3P”, in the absence of bound PI3P, non-symmetric hydrogen bonds are not present, and additional symmetric ß-Bridges are formed from anti-parallel stretches oriented towards the membrane. In simulation, Mol A switches to ß-Bridges instead of ß-sheets for residues indicated in Figure 2. All other hydrogen bonds are symmetric and not listed for the reversed Mol B to Mol A

| Dimer Intermolecular Hydrogen Bonds |  |  |  |
| --- | --- | --- | --- |
| Residue Mol A | Residue Mol B | Atomic Interaction | Backbone |
| Asp 30 | Ser 73 | OH—O |  |
| Tyr 71 | Gln 108 | O—HN |  |
| Tyr 71 | Gln 79 | O—HN | X |
| Ser 73 | Ala 29 | OH—O | X |
| Val 77 | Gln 79 | NH—O | X |
| Gly 82 | Tyr 107 | NH—O | X |
| Gly 83 | Arg 111 | NH—O | X |
| Leu 84 | Ser 112 | NH—O | X |
| Tyr 96 | Gln 108 | OH—O | X |
| Lys 97 | Gln 113 | NH—O |  |
| Lys 97 | Glu 114 | NH—O |  |
| Tyr 103 | Gln 108 | OH—O |  |
| Lys 105 | Gln 113 | NH—O |  |
| Non-symmetric |  |  |  |
| Glu 81 | Ser 73 | O—HO |  |

**Supplemental Movie 1: Monomer Steered Simulation**

Example steered simulation from one run.

**Supplemental Movie 2: Dimeric Steered Simulation**

Example initial steered simulation. With 1000 Kj restraint, the IP3P remains attached and is pulled from the membrane. These simulations were repeated with 2000 Kj restraint in order to calculate affinity for the ligand.

